# NetSyn: prokaryotic genomic context exploration of protein families

**DOI:** 10.1101/2023.02.15.528638

**Authors:** Mark Stam, Jordan Langlois, Céline Chevalier, Jean Mainguy, Guillaume Reboul, Karine Bastard, Claudine Médigue, David Vallenet

**Author notes:** Corresponding Author: Mark Stam.

## Abstract

**Background:** The growing availability of large prokaryotic genomic datasets presents an opportunity to discover new metabolic pathways and enzymatic reactions useful for industrial or synthetic biological applications. Efforts to identify new enzyme functions in this vast number of sequences cannot be achieved without bioinformatics tools and the development of new strategies. Standard methods for assigning a biological function to a gene are based on sequence similarity. However, complementary approaches rely on mining databases to identify conserved gene clusters (e.g., syntenies). In prokaryotic genomes, genes involved in the same pathway are frequently encoded in a single locus with an operonic organisation. This genomic context conservation is considered as a reliable indicator of functional relationships, and is therefore a promising approach for improving gene function prediction.

**Methods:** Here we present NetSyn (Network Synteny), a tool to group protein sequences based on the conservation of their genomic context rather than solely on sequence similarity. From a list of protein sequence identifiers, NetSyn searches for the corresponding genome entries to retrieve neighbouring genes. Corresponding protein sequences are grouped into families to define homology relationships and compute a synteny conservation score between the different extracted genomic contexts. A network is then created in which the nodes represent the input proteins and the edges indicate that two proteins share a conserved synteny. Finally, the network is partitioned into clusters grouping proteins with similar genomic contexts, using a community detection algorithm.

**Results:** As a proof of concept, we used NetSyn on two different datasets. The first one is the BKACE protein family (formerly named DUF849) which has previously been divided into isofunctional sub-families. NetSyn was able to go a step further by providing additional sub-families beyond those already described. The second dataset corresponds to a set of non-homologous proteins belonging to three different glycoside hydrolase (GH) families. These GHs are known to work cooperatively in Polysaccharide-Utilization Loci (PUL) and are therefore grouped together in the same genomic contexts. NetSyn was able to identify a locus grouping 3 GHs, involved in the degradation of xyloglucan, in 162 prokaryotic genomes.

**Discussion:** By highlighting conserved synteny in distantly related prokaryotic species, NetSyn enables functional links between proteins to be established beyond sequence similarity alone. We showed that NetSyn is efficient for exploring large prokaryotic protein families, enabling the definition of isofunctional groups and the identification of functional interactions between non-homologous enzymes. These features enable the prediction of new genomic structures that have not yet been experimentally characterised.

Finally, NetSyn is also useful for pinpointing annotation errors that have been propagated across databases, and for suggesting annotations for proteins lacking functional prediction. NetSyn is freely available at https://github.com/labgem/netsyn.

## Introduction

Advances in genome sequencing technologies have created an ever-increasing gap between the availability of protein sequences and the slow progress in their functional characterization and annotation (Galperin and Koonin, 2010). It is currently estimated that 23% of the families defined in the PFAM database are annotated as “unknown function” (Mistry et al., 2021).

This issue is further compounded by automatic annotation procedures, particularly those that rely solely on sequence similarity, which still exhibit a high false-positive rate, leading to annotation error rates of up to 80% for certain protein families (Goudey et al., 2022; Rembeza and Engqvist, 2021; Schnoes et al., 2009; Steinegger and Salzberg, 2020). Conversely, many known enzymatic activities have not yet been associated with protein sequences. In their review, Sorokina *et al*. show that more than 1000 EC numbers, referred to as orphan reactions, are not linked to a specific gene (Sorokina et al., 2014). To overcome the limitations of these methods, innovative strategies have been developed such as a combination of genome analysis and metabolic information (Chen et al., 2011; Kharchenko et al., 2006; Smith et al., 2012; Yamanishi et al., 2007), an integrative approach associating sequence family classification and controlled vocabulary (Jung et al., 2014), a Protein-Protein Interaction (PPI) network (Cozzetto et al., 2013; Peng et al., 2014; Zhao et al., 2016), and more recently, a machine learning technique based on natural language processing methods (Ofer et al., 2021).

One way to address the challenge of annotating gene function is to use genomic context information, which is considered a reliable indicator of functional relationships (Huynen and Snel, 2000; Janga et al., 2005; McClean et al., 2010; Rogozin et al., 2002) and has been used to support functional annotation (Ferrer et al., 2010; Lee et al., 2016; Vallenet et al., 2006). Genomic context methods can be classified into four categories (Ferrer et al., 2010): gene cluster, phylogenetic profile, gene fusion (or Rosetta Stone) and gene neighbour. The gene cluster method (or conserved synteny) is defined as the conservation of chromosomal proximity between genes (Overbeek et al., 1999; Tamames, 2001). Based on conserved genes in several organisms, the cluster method computes distances, in number of bases, between two adjacent genes transcribed in the same direction (Bowers et al., 2004; Overbeek et al., 1999). Small distances are considered to be a good indicator of co-regulated gene groups, as observed in operon structure (Brouwer et al., 2008) or Polysaccharide Utilisation Loci (PUL) (Bjursell et al., 2006). The phylogenetic profile method (Pellegrini et al., 1999) is based on the comparison of proteomes across genomes: genes A and B are functionally related if orthologs of A and B show similar patterns of presence and absence across many genomes. The gene fusion method (Marcotte et al., 1999) is based on the gene fusion events that can occur in some genomes. Pairs of proteins linked by such a fusion event are more likely to be functionally related and involved in the same biological process, for example, within the same metabolic pathway. The gene neighbour method (Bowers et al., 2004) calculates the distance between two genes in different genomes, based on the number of genes that appear between the two homologs plus one. Genes A and B are functionally related if orthologs of A and B across many organisms are located in close proximity on the chromosome (also called gene neighbours). Careful selection of genomes is required to avoid taxonomic bias (e.g., organisms from the same genus).

Further efforts have been made to incorporate gene context information into integrative approaches. For instance, the STRING database (Szklarczyk et al., 2017) stores known and predicted protein-protein interactions from different resources. These interactions can be physical (e.g., heterodimer) or indirect (e.g., gene regulation, signal relay mechanism) and are represented as a graph in which nodes correspond to proteins and edges to interactions. Such networks capture interactions between different protein families. However, the analyses are limited by the genomes included in the database and cannot support the exploration of an entire protein family. Other tools have been developed to search for conserved genomic contexts. For instance, Cblaster (Gilchrist et al., 2020) and CAGECAT (van den Belt et al., 2023), use the protein sequences of target genes and their genomic context as input data to perform a BLAST search in the NCBI reference genome database. Only blast hits separated by a maximum distance defined by the user (in bp) and belonging to the same organism and scaffold are retained. Another example is given by GCsnap (Pereira, 2021), which retrieves the genomic contexts by searching for the flanking genes of the input sequences; then it compares the retrieved genomic context. The Enzyme Function Initiative (EFI) (Oberg et al., 2023; Zallot et al., 2019) has also led to the development of EFI-GNT, a method based on the Genomic Neighbourhood Network (GNN), which enables the analysis of hundreds of genomes and the generation of large-scale datasets. It computes a Sequence Similarity Network (SSN) of the input protein sequences, in which groups of highly connected nodes are considered as a cluster. Then, neighbouring genes of each protein are retrieved from the European Nucleotide Archive (ENA) (Yuan et al., 2024) and considered related if they belong to the same PFAM family (Mistry et al., 2021). These methods do not rely on the creation of a true genomic context network, compared to a recent method called Syntenet (Almeida-Silva et al., 2023; Tao, 2025; Zhao and Schranz, 2017). Starting from whole-genome protein sequences, it searches for blocks of up to 50 genes conserved between two genomes and calculates a score for each of these blocks. A network is then created where the nodes represent genes in syntenic blocks and the edges represent syntenic relationships between genes. This network is divided using community detection algorithms and can be used to highlight synteny relationships. Syntenet is able to infer and analyse micro-synteny networks. It has been successfully applied on eukaryotic genomes (Guo et al., 2025; Li et al., 2025; Zhao and Schranz, 2019).

A strategy based partly on the search for conserved genomic contexts has enabled the division of the DUF849 family into isofunctional subgroups (Bastard et al., 2014). These results led us to develop NetSyn (Network Synteny), a tool able to detect conserved syntenies within distantly related organisms. Designed to analyse families from prokaryotic genomes, it links proteins into a network based on the conservation of their genomic context, highlighting specific loci involved in the same biological process. NetSyn provides several features to explore the synteny network, such as sequence clustering, visualisation of the genomic context of each input sequence, and integration of data from various resources. Overall, our tool quantifies the conservation of the genomic context of proteins from different prokaryotic genomes.

Compared to the tools mentioned above, NetSyn focuses on the genomic context of members of the analysed protein families to calculate a synteny score between their genomic context. It has been validated on two types of protein datasets. The first one is the initial set of sequences used for the analysis of the *β*-keto acid cleavage enzyme (BKACE) family (Bastard et al., 2014). The second one consists of sequences homologous to the three glycoside hydrolases present in a PUL that degrades xyloglucan. Overall, NetSyn is based on the principle that proteins sharing a similar genomic context across distantly related bacterial genomes are more likely to share similar functions or be involved in the same biological function (e.g., a metabolic pathway). Consequently, our method is well suited for partitioning diverse prokaryotic protein families into isofunctional clusters.

## Materials & Methods

### NetSyn workflow

NetSyn is divided into four steps (Fig.1): i) Genomic context extraction, ii) Protein family computation, iii) Synteny computation and scoring, iv) Synteny network computation and clustering (Supplementary data S1 illustrates how the computation time of NetSyn scales with the number of input sequences).

**Figure 1:**
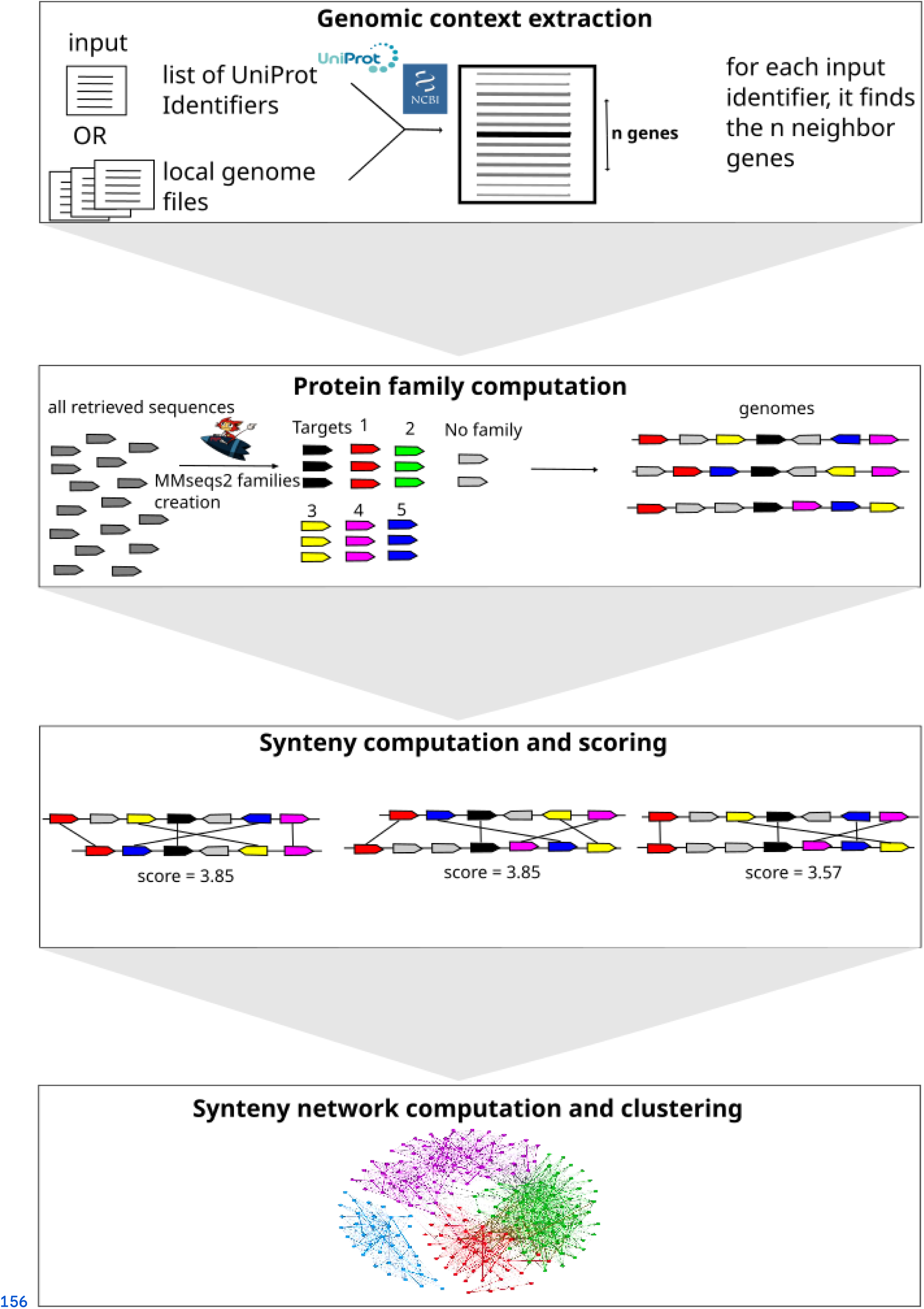
NetSyn Workflow. From a list of UniProt accessions, NetSyn downloads the corresponding genome assembly files or the user can provide their own assembly files. NetSyn then parses these files to recover the protein sequences of the target genes and their neighbouring genes (panel A). The retrieved protein sequences are clustered to define protein families (default values: 30% of identity on 80% of coverage) (panel B). Synteny relationships and scores are then computed for each pair of target proteins (panel C). A network is generated in which target sequences are represented as nodes and edges are created when a synteny with a score above a threshold (3 by default) is found. Finally, the network is partitioned using four different community detection algorithms (panel D).

#### Genomic context extraction

NetSyn takes as input a list of UniProt accession numbers (hereafter referred to as target proteins). The workflow uses the UniProt rest API to retrieve, for each entry, the corresponding nucleotide accession number and to download the associated genome assembly file (hereafter called the genome file) from the NCBI. Alternatively, NetSyn can accept local genome files listed in a tab-delimited file containing the following mandatory fields:

- **protein_AC**: target protein accession number in the genome file
- **protein_AC_field**: name of the field containing the protein_AC in the genome file
- **nucleic_AC**: nucleotide sequence accession number of the genome file
- **nucleic_File_Format**: format of the genome file (EMBL or Genbank)
- **nucleic_File_Path**: path to the locally stored genome file

Moreover, a metadata file containing attributes of the target proteins (e.g., a PFAM or INTERPRO family classification, the taxonomy of the corresponding organism) can be provided. These attributes can be displayed on the final synteny network (see below).

From the genome files, the sequence of the target proteins and their genomic context are extracted. The number of neighbouring genes to include can be adjusted according to the desired context size; by default, this value is set to 11 (i.e., 5 genes upstream and 5 genes downstream of the target gene). As the average number of genes in prokaryotic operons ranges from 2 to 3, this window size is generally sufficient to detect most conserved gene clusters (Zheng et al., 2002).

NetSyn also retrieves the taxonomic lineage of the organisms from which the input sequences originate by querying the NCBI API using the taxonomic identifier extracted from the genome file. As a result, the taxonomic lineage provided by NetSyn is derived from the NCBI and is regularly curated (Schoch et al., 2020). This taxonomic information can be mapped onto the final network and can also be used to remove redundancy (see “Merging nodes and network reduction” section).

#### Protein family computation

The purpose of this step is to define homology relationships between proteins. All the retrieved proteins (target proteins and proteins from neighbouring genes) are clustered using MMseqs2 (Steinegger and Söding, 2017). A common rule of thumb is that proteins sharing at least 30% sequence identity over at least 80% of their length are likely derived from a common ancestor (Rost, 1999). We used these as default values (these parameters can be modified). The resulting clusters enable NetSyn to define a set of homologous protein families which are found in the context of the target proteins. These families are then used to compute synteny conservation for each pair of target proteins (Fig. 1).

#### Synteny computation and scoring

Genomic context conservation between two target proteins is computed using an exact graph-theoretical approach (Boyer et al., 2005). From the two genomes containing the target protein genes (denoted T_A_ and T_B_), two networks (N_A_ and N_B_) are constructed containing the target protein genes and their neighbouring genes as vertices (G_A1-n_ and G_B1-n_, according to the window size parameter), with edges representing their colocalization along the genome. To account for potential gaps (i.e. genes within a conserved synteny that lack homologs in the other genomes), a partial transitive closure is applied to N_A_ and N_B_, linking genes separated by no more than n genes (synteny gap parameter, set to 3 by default). Edges are then added between N_A_ and N_B_ to represent homology relationships between genes belonging to the same protein families computed previously. Finally, a common connected component (CCC) between the two networks is identified by constructing a multigraph in which vertices from N_A_ and N_B_ are merged based on homology relationships. These CCCs correspond to conserved synteny groups. Figure 2 illustrates the method, showing the extracted genomic contexts around T_A_ and T_B_ (panel A) and the resulting detected synteny (panel B). G_A1_ and G_B2_ genes are not part of the conserved synteny because the number of genes between G_A1_ and T_A_ in genome A exceeds the gap parameter threshold.

**Figure 2:**
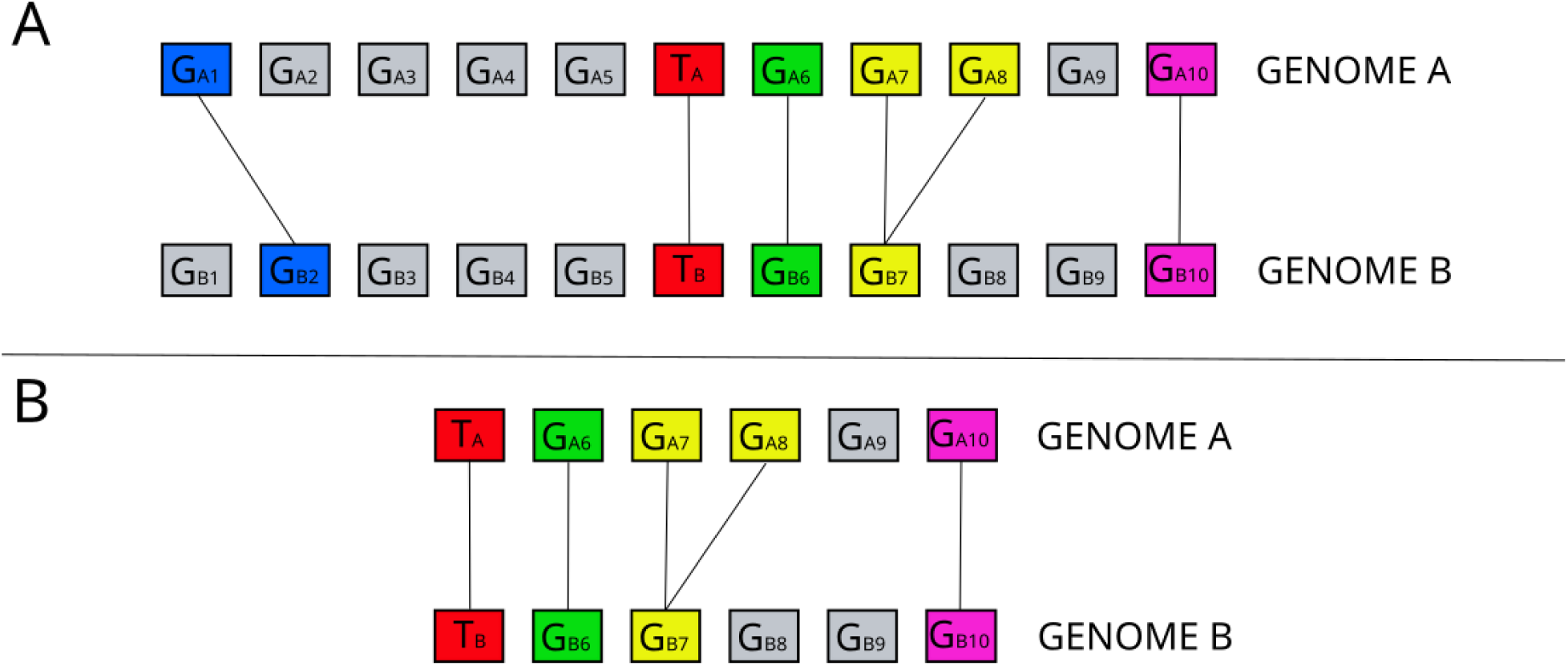
Example of synteny computation between two target protein genes. A) Genomic context extraction from two target protein genes: T_A_ from genome A and T_B_ from genome B. In this example the window size is set to 11, meaning that NetSyn considers 5 genes upstream and 5 genes downstream of each target gene to extract the genomic contexts in both genomes. Homologous genes between the two genomic contexts are represented by coloured rectangles and linked by edges. B) Detected synteny. The synteny predicted by NetSyn includes the following pairs of homologous genes: T_A_/T_B_, G_A6_/G_B6_, G_A7_/G_B7_, G_A8_/G_B7_, G_A10_/G_B10_. The G_A1_/G_B2_ homologs are not part of the conserved synteny because the number of genes between G_A1_ and T_A_ in genome A (i.e., 4 genes) is greater than the gap parameter threshold (set to 3 by default). 9 genes are conserved in the synteny out of 12 genes, including 3 gap genes (G_A9_, G_B8_ and G_B9_). The Synteny Score is thus equal to (9/2)*(9/12)=3.375.

Once a conserved synteny has been identified for a pair of target proteins, a score is calculated as follows:

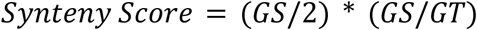

where GS is the number of genes involved in the synteny group, and GT is the total number of genes in both genomic contexts, including gap genes. This score represents the average number of genes in the synteny, penalized by the presence of gap genes. For example, in Figure 2 (panel B), the genomic context contains 9 syntenic genes out of a total of 12 genes, including 3 gap genes (G_A9_, G_B8_ and G_B9_). Therefore the Synteny Score (SS) is equal to (9/2) * (9/12) = 3.375.

#### Synteny network computation and clustering

Once all syntenies and scores have been computed between target proteins, NetSyn builds a synteny network in which nodes represent the target proteins. Two target proteins are connected by an edge if a conserved synteny is detected between them. The weights of edges are equal to synteny scores, highlighting the most conserved syntenies. To retain only informative syntenies, NetSyn applies a default threshold of 3 on the synteny score, although this value can be adjusted by the user. Target proteins that are not connected to any other target protein are excluded from the final network.

To group target proteins sharing similar genomic contexts, several community detection algorithms are implemented in the NetSyn workflow: MCL, Walktrap, Louvain and Infomap. The Markov Cluster algorithm (Dongen, 2000) (MCL) is an unsupervised algorithm based on the simulation of flow in graphs (it simulates random walks within a graph). The Walktrap algorithm (Pons and Latapy, 2005) is also based on random walks and performs well on large networks (> 1000 vertices). The Louvain algorithm (Blondel et al., 2008) is based on the comparison of the edge density inside and outside a community (i.e., a cluster). The Infomap algorithm (Rosvall and Bergstrom, 2008) tries to minimise the length of a randomised walk trajectory into a cluster.

These four algorithms are independently applied to identify groups of target proteins sharing similar genomic contexts. In the synteny network, the resulting clusters are visualized using distinct colors, with each color representing a group of related target proteins. By default, the clustering method producing the smallest number of clusters is displayed in the HTML output. Our analyses indicate that this method frequently corresponded to the one that minimized the formation of ‘small’ and non-pertinent clusters (i.e., clusters containing between 2 and 5 nodes). However, users can also explore the results obtained with other clustering methods, both in the HTML output and in network visualisation software.

#### Cluster quality

In the clustering framework based on genomic context conservation, a cluster is considered coherent when most nodes within the cluster are interconnected. To assess this property, we use the alpha index which gives a measure of connectivity within a given cluster. The alpha index is the ratio between the number of cycles (i.e., closed paths that start and end at the same node and pass through nodes only once) and the estimated total number of cycles in the network. It is calculated using the following formula: alpha = (e-v)/((v(v-1)/2)-(v-1)) where e is the number of edges and v is the number of vertices. A value of 1 indicates a fully connected cluster, while a value of 0 means a completely disconnected cluster (i.e., a minimally connected cluster without cycles).

### NetSyn Output

We provide two output files for exploring and analysing the network: a graphML file which can be opened with network visualisation tools such as Gephi or Cytoscape, and an HTML file that can be viewed in a Web browser (Fig. 3). In the graphML output, information from the metadata file and the extracted taxonomic lineage are mapped onto the network.

**Figure 3:**
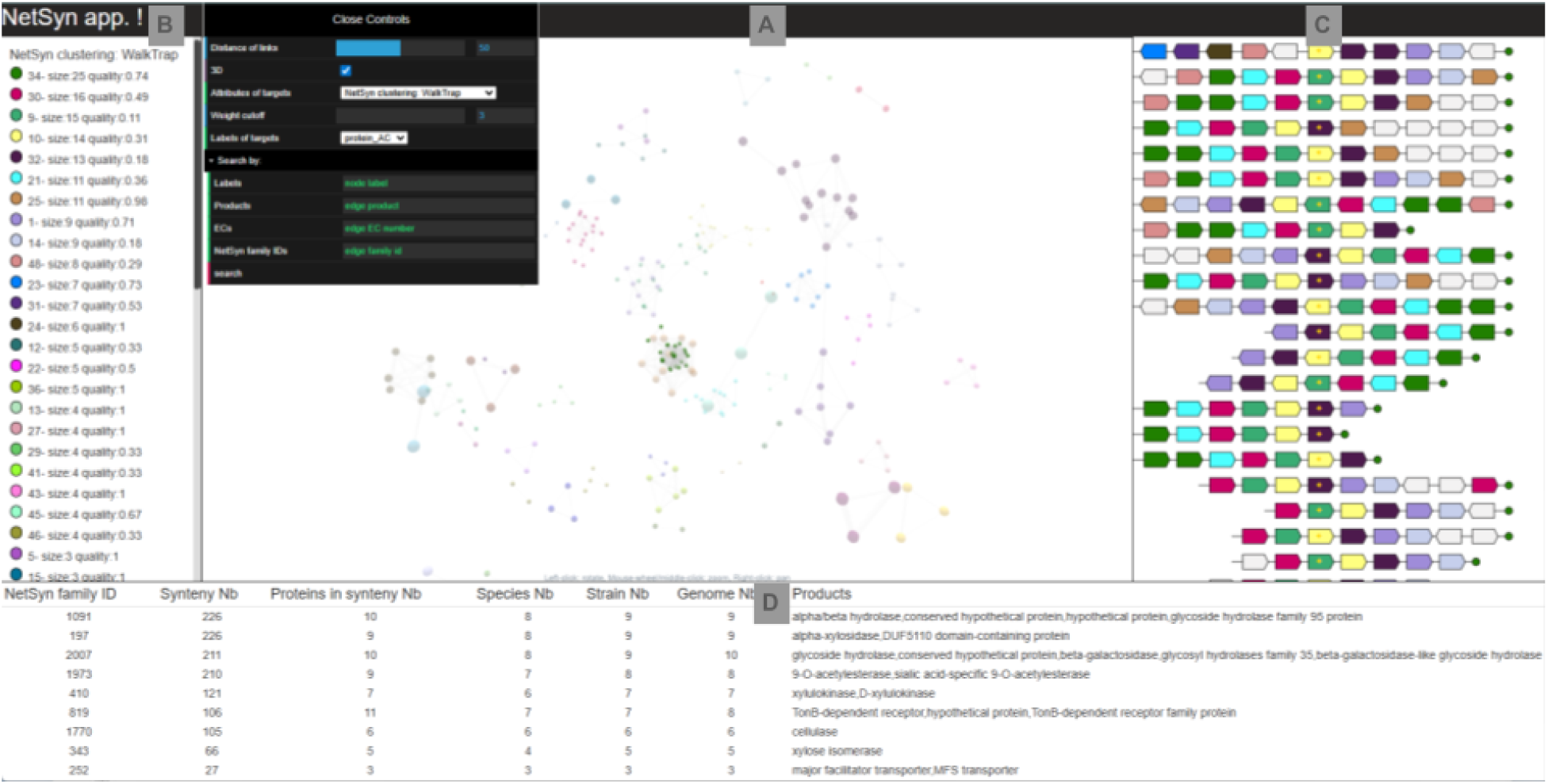
Visualisation of the NetSyn results. Panel A displays the NetSyn network. The control panel allows customization of node colors based on the “Attributes of targets” selection, which includes NetSyn clusters (obtained with one of the four available algorithms), taxonomic ranks, or user-defined metadata. Nodes can be selected using the “Search by” functionality, with the following search fields: Labels (target gene names), Products (functional descriptions), ECs (annotated EC numbers), NetSyn family IDs (protein family identifiers). In this panel, the network can be zoomed in or out and rotated. Panel B provides the color legend for the currently selected ’Attributes of targets’ ordered by decreasing number of target proteins. The user can click on a specific coloured item (e.g., cluster 1 obtained with the walktrap method) to highlight the corresponding nodes in the network. Panel C displays the genomic context of the selected cluster. In this view, each arrow represents a gene and homologous genes (i.e., genes belonging to the same protein family) share the same color. Target genes are marked with a golden star. When the user hovers over a gene, a pop-up window displays the following information: protein family identifier, strain name, UniProt accession, locus tag, product, gene name, and EC number (if available). Panel D displays a summary table of the protein families identified within the selected cluster. The table includes the following columns: NetSyn family ID (protein family identifier), Synteny Nb (number of syntenies involving the protein family), Proteins in Synteny Nb (number of proteins of the family involved in a synteny), Species Nb (number of distinct species), Strain Nb (number of strains), Genome Nb (number of genomes), Products (concatenated protein annotations of the family ordered by decreasing frequency).

The HTML view is divided into four panels. The synteny network is displayed in the central panel (A), where each node represents a target protein, and an edge between two nodes indicates that a conserved genomic context with a synteny score above a given threshold (3 by default) has been identified between them. Nodes can be colored according to different attributes, such as cluster identifiers, genome taxonomy or user-defined metadata. The control panel at the top of the A panel allows users to select the attributes and information displayed on the synteny network (see Fig. 3 captions). The left panel (B) displays the network legend, which depends on the parameters selected in the control panel, the identifier, the number of target proteins and the alpha index (see above) of each cluster (Fig. 3B). Selecting a coloured item in panel B, automatically highlights the corresponding nodes in the synteny network (panel A). In addition, the genomic contexts of the corresponding target proteins are displayed on the right panel (C). Each context is centered on the target protein gene, which is marked with a yellow star, surrounded by neighbouring genes belonging to the conserved synteny. In this view, homologous genes share the same colour (*i.e*., they belong to the same protein family). The bottom panel (D) provides, for the selected group of target proteins (panel B), a summary table of the protein families found within the conserved syntenies, including the number of proteins, the number of corresponding species, strains and genomes, as well as functional annotations.

Two additional files are also created:

- A <OUTPUT directory name<_Report_1.txt file, which summarizes the results of the different steps (e.g., number of target proteins without corresponding Genbank entry or assembly file, number of clusters obtained with each clustering method, etc.).
- A <OUTPUT dir name<_Settings_1.yaml file, which records the parameters used to run NetSyn.

#### Merging nodes and network reduction

In some cases, the resulting network can be large and difficult to visualise. To address this issue, NetSyn provides a node-merging option to reduce network complexity. When this mode is enabled, users must select a clustering method and a property on which nodes will be merged (i.e., a metadata column or a taxonomic rank retrieved by NetSyn). NetSyn first performs a standard analysis by identifying synteny clusters among all input target proteins. It then merges nodes belonging to the same NetSyn cluster and sharing the selected property into a single node in the output graph. This approach can be used, when necessary, to reduce the impact of taxonomic redundancy on network visualization. (see Supplementary data S2 for an example)

## Results & Discussion

The NetSyn method was evaluated on two datasets. The first dataset consists of a family of homologous proteins, the *β*-keto acid cleavage enzymes family, which has been well studied with experimental validations. The second dataset comprises a mixture of three evolutionarily distinct protein families that are known to be co-localized and to interact in the degradation of xyloglucan.

### Dividing a homologous family

The *β*-keto acid cleavage enzyme family (BKACE), initially called DUF849 (Domain of Unknown Function) in the PFAM database, has been extensively characterised using high-throughput enzymatic screening (Bastard et al., 2014). Activity profiles obtained from the screening of 124 representative BKACE sequences across 20 substrates showed a strong correlation with active-site profiles. At the time of the analysis, the BKACE family contained 725 sequences from 337 bacterial genomes (mostly from Pseudomonadota), as well as 3 archaeal, and 1 eukaryotic genome. This family was divided into seven main groups based on a combination of different clustering methods, including a structural classification of their active sites computed using the Active Sites Modeling and Clustering (ASMC) method (Bailly et al., 2025; de Melo-Minardi et al., 2010). The ASMC groups correlate well with the biochemical characterizations. In this study, we aimed to compare ASMC and NetSyn clusters to assess the extent of overlap between the two methods.

We retrieved the identifiers of the 725 sequences used in the ASMC analysis and submitted them to NetSyn. Of the 725 entries, 159 could not be associated with a Genbank assembly file because the corresponding UniProt entries were obsolete. Among the remaining 566 entries, 86 lacked a relevant conserved genomic context (i.e., with a synteny score >=3). Therefore, 480 input entries were retained in the final network. The Louvain algorithm (see ‘Synteny network computation and clustering’ section) led to a minimum number of clusters (33), with cluster sizes ranging from 2 to 68 sequences (Table 1). Most of these clusters show good agreement with the seven experimentally validated groups previously defined by the ASMC method. Although the overall partition similarity is moderate (Adjusted Rand Index = 0.41), the high homogeneity score (0.73) indicates that NetSyn consistently groups experimentally validated isofunctional proteins together while further subdividing them according to their conserved genomic contexts (Hubert and Arabie, 1985; Rosenberg and Hirschberg, 2007).

**Table 1:**
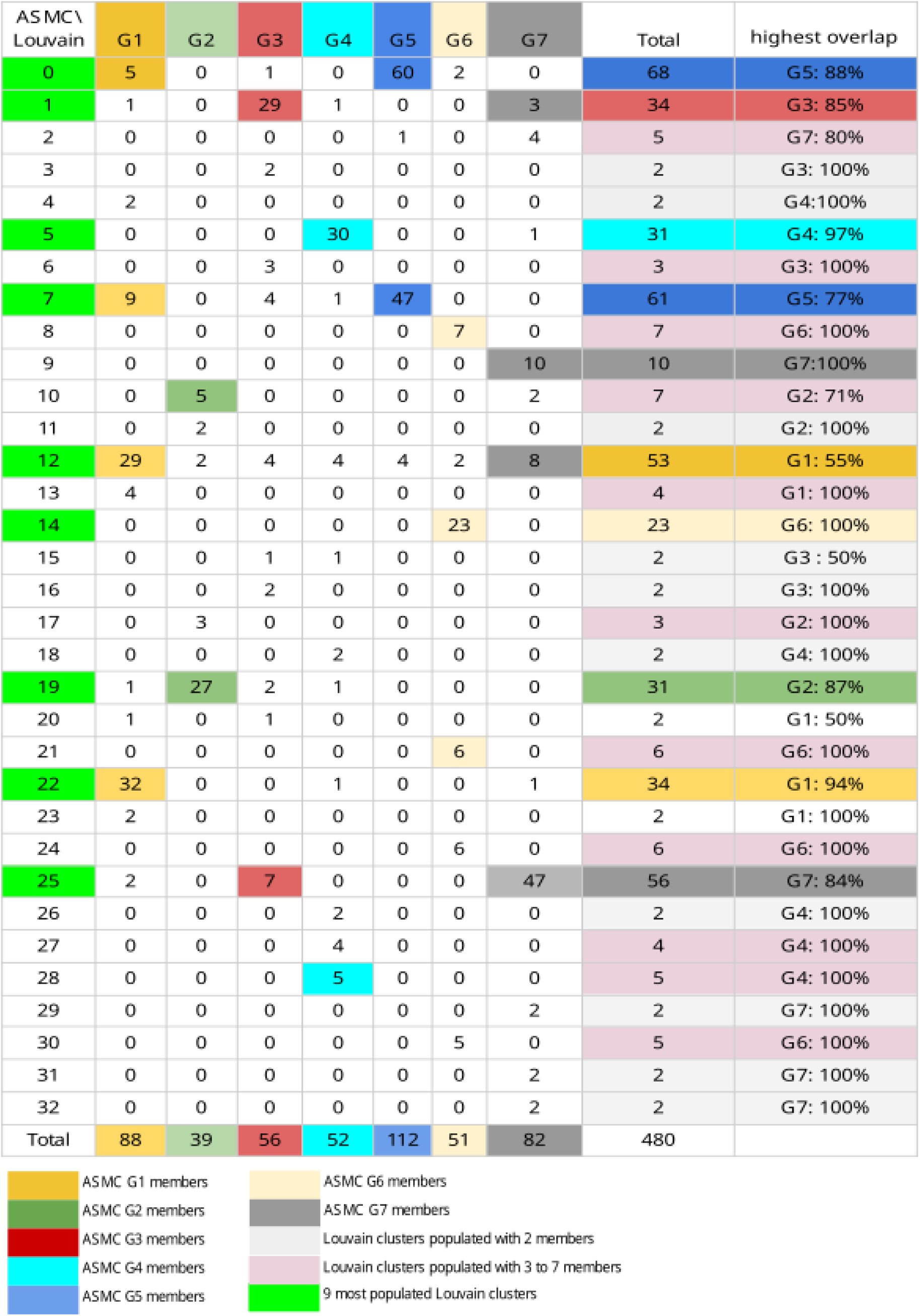
Comparison between ASMC groups and Louvain NetSyn clusters.

It should be noted that 12 clusters contain only 2 proteins. These clusters are considered as singleton and are of little interest for our analysis as they do not gather sufficient contextual genomic information to highlight a biological process. Most of the sequences (391) are clustered into 9 Louvain clusters (0,1,5,7,12,14,19,22,25) with a number of sequences ranging from 23 to 68. Remarkably, 89% of these clusters correlate well with the 7 ASMC groups (named here after G1 to G7; Table 1): between 77 to 100% of the members of one specific NetSyn cluster belong to the corresponding ASMC group. The only exception is cluster 12 in which only 55% of the sequences belong to the ASMC group G1. Indeed, the cluster number 12 contains sequences from all the groups defined by the ASMC method (Table 1). Visualisation of the network generated by NetSyn on BKACE data is available at https://doi.org/10.5281/zenodo.22081334. By design, NetSyn built clusters of proteins according to their genomic context, while ASMC groups them according to their active sites, which directly reflects the type of substrate they can catalyse (negative, positive, polar or hydrophobic substrates). The G7 group is the perfect example of this difference. Proteins lacking at least one residue of the catalytic active site were grouped together in the ASMC G7 group and were shown not to transform any keto-acids during enzymatic screening (Bastard et al., 2014): they have been annotated as non-BKACE enzymes. The G7 group is therefore a heterogeneous group whose functions could not be studied in our previous study (Bastard et al., 2014). Interestingly, the context of the proteins clustered by NetSyn provides sufficient clues to investigate new metabolic pathways in which G7 proteins could be involved (Table S1). The proteins from the ASMC G7 group have a large diversity of genomic contexts and have thus been distributed into 4 large Louvain clusters (Table 1). More than half of the enzymes found in group G7 are also found in the Louvain cluster 25, which includes a locus containing a BKACE-like enzyme, a quinone oxidoreductase, a taurine dioxygenase, an MSF transporter, a serine hydrolase and a transcriptional regulator (Supplementary data S3). This metabolic cluster has been described in *Rhodococcus erythropolis* (Actinomycetes class) in the STRING database (Szklarczyk et al., 2023) but further experimental studies are needed to determine whether this is a metabolic pathway.

The ASMC G1 group was split into two highly populated NetSyn clusters (clusters 12 and 22; Table 1). Examination of the genomic context of cluster 12 shows that it is involved in the conversion of valine to leucine, while cluster 22 is probably involved in a pathway that remains to be discovered from annotations of neighbouring genes (Supplementary data S3). The G5 group was also divided into two highly populated NetSyn clusters (clusters 0 and 7). Their genomic contexts outline the two pathways involved in carnitine degradation (Supplementary data S3).

The advantage of NetSyn over ASMC is that it is not dependent on a molecular modelling threshold. For example, seven proteins have been classified by the ASMC method in the G3 group (for which there is no specific substrate signature), because their 3D models contain all the catalytic residues. However, according to their metabolic neighbours, these proteins have been classified in the NetSyn cluster 25 which contains sequences mainly from the G7 ASMC group (Table 1). The same applies to four proteins grouped in G1, G3 and G4, now classified in the Louvain cluster 19: the genomic neighbourhood of these four genes shares two genes (Acetoacetate:butyrate CoA-transferase *α* subunit and *β* subunit) with other members of cluster 19 (Supplementary data S4). Another example is given by 2 proteins, the 3D models of which were of poor quality (sequence identity with the reference structure respectively equal to 23% and 19%), that were grouped together, by default, in cluster G7. However, these proteins have been more coherently linked by NetSyn to putative metabolic functions present in their conserved genomic context (Table 1): probably fatty acid biosynthesis for protein C0ZQU4 (cluster 1) and deoxyribose catabolism for protein A5V766 (cluster 22).

NetSyn is also largely independent of the taxonomic ranks of the clustered proteins. For instance, cluster 19 contains proteins from organisms belonging to several distinct taxonomic classes, including Clostridia, Fusobacteriia, Bacteroidia, Betaproteobacteria, and Gammaproteobacteria. All the major NetSyn clusters (i.e., 12, 1, 0, 14 and 25) have a wide range of taxonomic classes. One exception is cluster 5 (31 members) which contains only Alphaproteobacteria, mainly from the order Rhodobacterales.However, its conservation may be functionally driven, as the genomic context contains genes involved in a metabolic pathway for the degradation of aromatic compounds (see Supplementary Data S3). To avoid biases caused by the presence of comparable, or even identical, genomic contexts of closely related organisms, users can apply the node-merging option of NetSyn. Comparing the NetSyn graphs obtained before and after applying the merge option, allows the user to identify which cluster is populated due to the diversity of the taxonomy.

Overall NetSyn was able to find ASMC groups and to suggest even more detailed clusters. As ASMC has been shown to reflect *in vitro* activities, the agreement between these two methods indicates that NetSyn is able to cluster iso-functional enzymes. NetSyn clustering is influenced more by the composition of genomic neighbourhoods than by the taxonomic origin of the target proteins, as most of the clusters characterised here are highly taxonomically diverse. Manual curation of the NetSyn clusters based on literature also has demonstrated that putative metabolic pathways can be inferred.

### Interconnectivities of families in metabolic pathways

A major advantage of NetSyn is its ability to analyse non-homologous but functionally related genes. Indeed, a large proportion of genes in prokaryotes, and a smaller fraction in eukaryotes, are physically linked on chromosomes and involved in a specific biological function (Nützmann et al., 2018). The most widely studied genetic organisation is the operon (Jacob and Monod, 1961), in which genes are transcribed into a single polycistronic mRNA. Operons are often conserved across species through vertical inheritance, and are therefore relatively easy to predict.

Another example of gene organisation is the Polysaccharide Utilization Loci (PULs), which have been described in Bacteroidetes. These loci consist of physically linked genes involved in carbohydrate metabolism, including two key proteins composing the transport system: a TonB dependent receptor (namely SusC) and a carbohydrate binding protein (namely SusD) (Bjursell et al., 2006). Therefore, a strategy for PUL prediction is based on the detection of the susC/susD pair within a cluster of Carbohydrate Active Enzymes (CAZymes) (Terrapon et al 2015). However, the SusC/SusD paradigm has been challenged by the discovery of alternative sugar transport systems associated with clusters of CAZyme (Larsbrink et al 2014; O Sheridan et al, 2016), as well as by evidence that the SusC/SusD system is not necessarily co-localised with CAZyme genes (Ficko-Blean et al 2017). Therefore, methods such as NetSyn, which highlight conserved genomic contexts involving non-homologous genes, are required to reliably detect gene clusters such as PULs.

Xyloglucans (XyGs) constitute a family of ubiquitous and abundant plant cell wall polysaccharides. They are composed of (1->4)-β-glucan backbone substituted with *α*(1->6)-xylosyl side chains, which can be extended with additional glycosyl residues (Fig. 5 panel B) (Grondin et al 2017). Xyloglucan can be degraded by a complex gene locus in the saprophytic Gram-negative bacterium *Cellvibrio japonicus,* referred to as a XyGul (Xyloglucan Utilisation Loci). This locus is composed of three glycoside hydrolases (GHs) from the GH31, GH35 and GH95 families defined in the CAZy database (Drula et al., 2022) as well as a TonB-dependent receptor (TBDR) (Larsbrink et al 2014). This particular locus has also been detected in several *Xanthomonas* strains (Vieira et al., 2021).

This example aims to demonstrate that NetSyn can identify loci containing proteins from non-homologous families, without being limited to closely related organisms. We therefore used the three key genes (xyl31A, bgl35A and afc95A) from *Cellvibrio japonicus Ueda107* as reference genes to search for homologous proteins in the UniProt database using BLAST (default parameters) (Altschul et al., 1997). Homologous sequences were selected by choosing those with at least 30% identity and 80% coverage, resulting in a dataset of 15,452 candidates sequences (2,212 homologous to xyl31A, 1,756 homologous to bgl35A and 11,484 homologous to afc95A), from a wide range of different genomes (4,268 Bacteria, 27 Archaea and 842 Eukaryota).

This dataset was submitted to NetSyn. Of the 15,452 entries, 251 could not be associated with a GenBank identifier and 379 with an assembly file. In addition, no conserved genomic context was identified for 1481 entries. The final network, which gathers 13,341 entries, is available at: https://zenodo.org/records/22047144.

The Louvain algorithm produced 310 clusters including 6 clusters with sizes ranging from 910 to 3005 sequences, 7 clusters ranging from 106 to 509 sequences, 15 clusters ranging from 20 to 89 sequences and 282 clusters ranging from 2 to 18 sequences (figure 4A).

**Figure 4:**
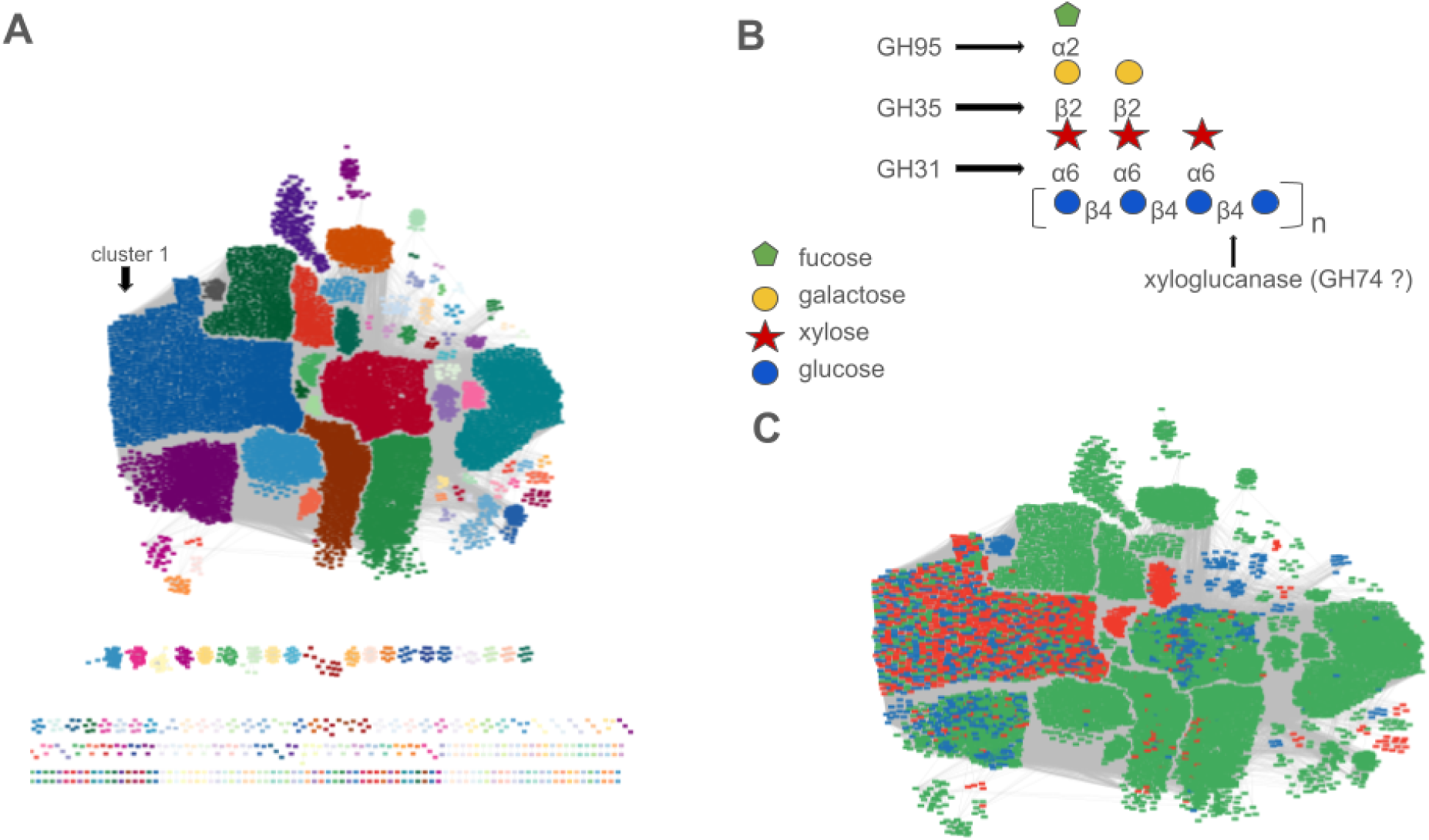
Panel A: Final NetSyn network visualized with Cytoscape. In the center of the figure is shown the biggest connected components of the network. Each cluster is represented by a given color. Panel B : Xyloglucan’s structure and enzymes evolving in the C. japonicus Xygul (from Grondin et al (Grondin et al., 2017)) Panel C : Zoom on part of the NetSyn network drawn by Cytoscape showing the biggest connected components. In each cluster, homologous proteins to GH31, GH35, and GH95 are respectively colored in blue, red and green.

Only six NetSyn clusters group sequences belonging to the three GH families (clusters 1, 7, 6, 5, 2 and 10), which are most often located in different loci. Among them, only cluster 1 brings together the three GH families (GH31, GH35 and GH95) from co-localized genes including the three reference GHs from *C. japonicus*. By investigating the loci retrieved in cluster 1, we identified 166 putative XyGuls in 162 distinct genomes belonging to different classes of Proteobacteria. Forty XyGul loci have previously been described in various *Xanthomonas* strains (Vieira et al., 2021), of which 19 were recovered by NetSyn. The remaining loci were not detected because the corresponding proteins were either not present in Uniprot, or had less than 30% identity with the sequences of *C. japonicus,* and were therefore excluded from the initial dataset.

Interestingly, although XyGULs have been described exclusively in Gammaproteobacteria, our analysis revealed the presence of similar loci in Betaproteobacteria and Alphaproteobacteria. A more in-depth analysis of the XyGULs identified in Alphaproteobacteria is required, as in two of them, the exo-*α*-1,2-L-fucosidase (EC 3.2.1.63) belonging to the GH95 family is absent, and appears to be replaced by a GH29 enzyme, a family that also contains exo-alpha-1,2-L-fucosidases (Wu et al., 2023). It is worth noting that the locus described in *Cellvibrio japonicus* lacks xyloglucanase activity (EC 3.2.1.151), which is found in the GH74 family, which is essential for the degradation of the xyloglucan backbone (Larsbrink et al., 2014; Vieira et al., 2021). Notably, among the 162 genomes identified in our analysis, 88 contain a XyGUL including a GH74 *(Xanthomonas, Massilia, Pseudoxanthomonas and Pseudoduganella)* (Fig. 5, Supplementary data S5). This supports the conclusion that we have detected loci involved in xyloglucan degradation.

**Figure 5:**
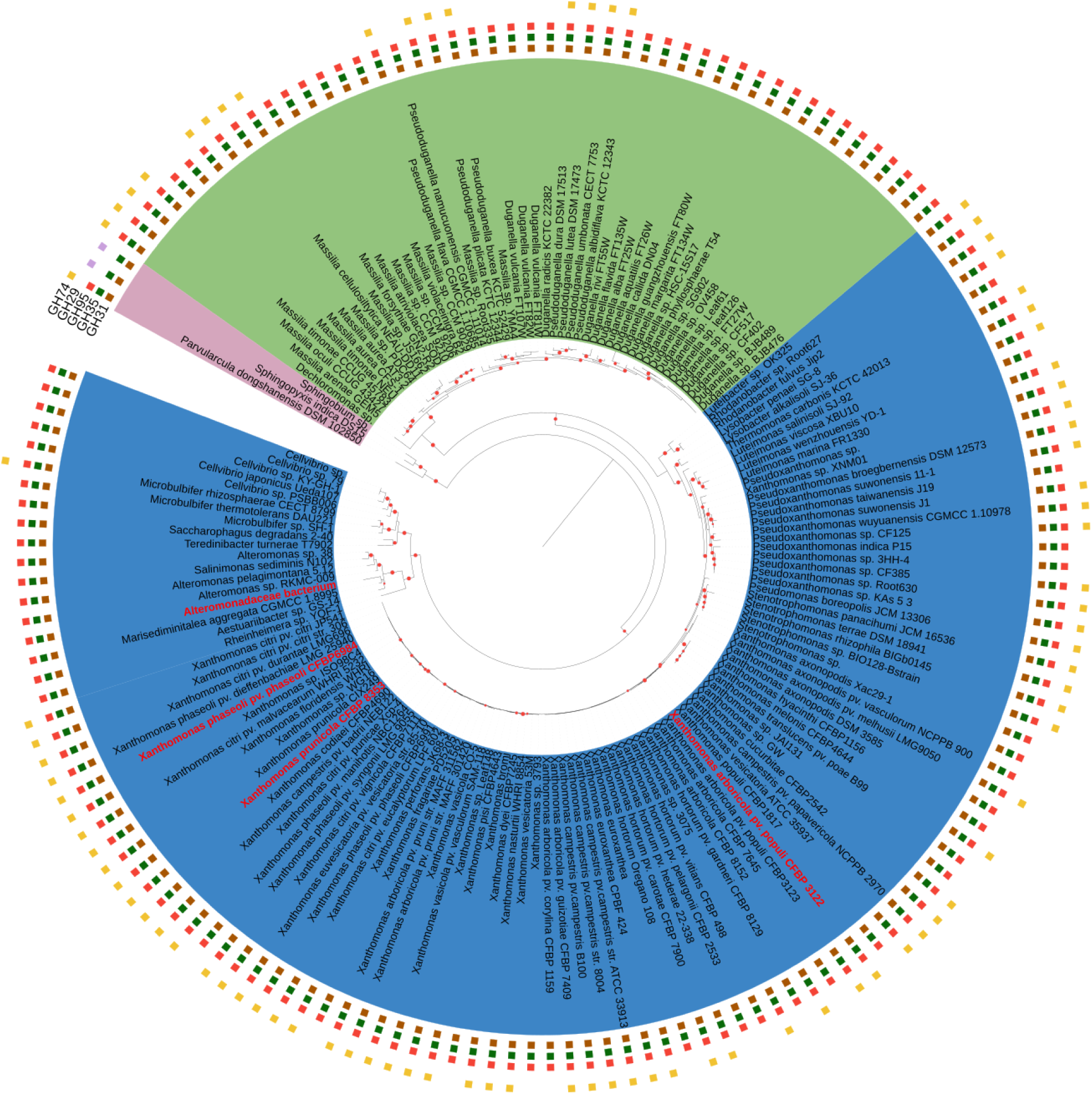
The tree was created using 16S rRNA sequences from organisms in which a XyGUL locus similar to that of C. japonicus was identified. From the center to the periphery, the first layer displays organism names: with each genus represented by a specific background colour (Alpharoteobacteria in pink, Betaproteobacteria in green and Gammaproteobacteria in blue). The second layer shows the presence of GH31 (brown squares), the third GH35 (green squares), the fourth GH95 (red squares), the fifth GH29 (purple squares) and the outermost layer GH74 (yellow squares). Bootstrap values above 80 are indicated with a red circle.

Moreover, NetSyn unveils other genes that are involved in xylose metabolism (or more generally in its degradation), such as xylulokinase and acetylesterase. For instance, specific transporters (ABC sugar transporters, ABC xylose transporters) and enzymes (D-xylose-1-dehydrogenase) were found within the putative XyGUL loci previously identified (Stephens et al., 2007). These results indicate that the locus detected in the 160 organisms, gathered in NetSyn cluster 1, is dedicated to xyloglucan degradation. Indeed, organisms in which a xyloglucan degradation locus has been predicted by NetSyn belong to different classes: Alphaproteobacteria, Betaproteobacteria and Gammaproteobacteria (Fig. 5). This observation suggests that the conserved synteny within this NetSyn cluster is due more to a functional conservation than to taxonomic conservation.

The use of NetSyn enabled the detection of the XyGUL locus in many different phyla, including Alphaproteobacteria (Table 2). Finally, we show that NetSyn can group together enzymes acting synergistically in the same degradation system. However, experimental characterisation is still required to validate these predictions.

**Table 2:** Comparison of tools for exploring genomic context.

|  | NetSyn | Syntenet (R package)/<br>SynNet-Pipeline<br>(bash and R script) | EFI-GNT | GCsnap | CAGECAT |
| --- | --- | --- | --- | --- | --- |
| Purpose | Partition prokaryotic protein families based on synteny conservation | Infer synteny networks from whole-genome protein sequence data | Analyse genomic contexts of protein families from sequence similarity network (SSN) clusters | Compare genomic contexts from a list of proteins | Search for homologous gene clusters |
| Input | List of proteins from UniProtKB or GenBank/ENA files | Complete genomes in GFF and CDS/protein in FASTA formats | EFI-EST sequence similarity network | List of proteins from NCBI Entrez, UniprotKB, UniRef, GeneID or Ensembl | GenPept protein accessions, GenBank gene clusters, Pfam identifiers, CDS/proteins fasta file or GenBank file with regions of interest |
| Output | Network of proteins clustered by synteny conservation | Network of genes clustered by syntenic relationships and phylogenomic profiles | Network of SSN clusters linked to PFAM domains | Genomic map showing synteny conservation | Gene presence/absence matrix and genomic map of synteny conservation |
| Output format | Tsv files, interactive html graph and graphML file | Text files, tables and graphical outputs | XGMML files and SQLite databases | Text files, interactive HTML file | Interactive HTML file |
| Type of pipeline | Python pipeline; fully automatic | R and bash scripts. Genome file download and protein family member selection are manual while homology search, synteny computation, network construction and infomap clustering are automated | Web server | Python pipeline, genome Download (automatic or manual) and homology search (automatic) | Web server |

### NetSyn limitation and comparison with other tools

One limitation of NetSyn is that clustering may be biased by the selection of sequences from organisms belonging to a narrow taxonomic range (e.g., organisms from the same species or genus). Indeed, a lack of genomic diversity in the input dataset may lead to the detection of synteny clusters driven primarily by phylogenetic relatedness rather than functional conservation. To overcome this problem, we propose reducing redundancy by merging nodes within NetSyn clusters that share the same taxonomic rank or metadata (see ‘Merging nodes and network reduction’ section). This approach simplifies network visualization by reducing the visual prominence of clusters grouping sequences from closely related organisms, without modifying the initial clustering. The simplified network can then guide the selection of a subset of target proteins that best represents the taxonomic and genomic context diversity of the dataset before rerunning NetSyn, thereby potentially improving clustering. Because target proteins from genomes belonging to the same species or genus often share similar genomic organisation, we recommend merging nodes at least at the genus level (see Supplementary data S2 as an example). However, no single taxonomic rank can be recommended universally. Even within a single species, homologous target proteins may have distinct genomic contexts, and thus potentially different functions, as a result of independent horizontal gene transfer events. Conversely, redundancy may also occur at higher taxonomic levels in some datasets.

Another current limitation concerns the number of sequences that NetSyn can handle: beyond 10,000 sequences, graphML files may be too large to be opened by Gephi or Cytoscape, and the generated HTML file may exceed browser size limits. Furthermore, although NetSyn can process proteins from eukaryotic organisms, it was primarily designed for prokaryotic protein families and may therefore be affected by features specific to eukaryotic genomes, such as large tandem gene arrays.

Several tools can perform synteny analysis (e.g., Syntenet, EFI, GCsnap and CAGECAT), but direct comparison with NetSyn is challenging because these tools differ substantially in their objectives and methodologies. For instance, NetSyn was developed to explore prokaryotic protein families in order to uncover functional relationships in distantly related bacteria, whereas Syntenet focuses on whole-eukaryotic-genome comparisons, notably to support phylogenetic reconstruction (Almeida-Silva et al., 2023; Guo et al., 2025; Li et al., 2025; Tao, 2025; Zhao and Schranz, 2019, 2017).

EFI groups proteins based on sequence similarity rather than genomic context conservation. Although this approach is robust, it relies on Sequence Similarity Network (SSN) driven solely by sequence homology, without considering conservation of genomic context within the studied families. Consequently, EFI does not support analyses that combine proteins of distinct evolutionary origins, unlike NetSyn (Oberg et al., 2023).

GCsnap is methodologically similar to NetSyn, as it retrieves and compares the context of input sequences. However, NetSyn goes further by grouping protein sequences according to the similarity of their genomic context (Pereira, 2021).

Finally, CAGECAT enables the search for homologous gene clusters in the NCBI database, but this approach requires prior knowledge of the gene cluster composition. In contrast, NetSyn can identify conserved genomic contexts without any prior information about their composition (van den Belt et al., 2023) (cf. Table 2).

## Conclusion and perspectives

We presented NetSyn, a novel tool that links prokaryotic protein sequences within a network based on the conservation of their genomic context. It fills the gap left by current methods that rely solely on sequence similarity or active-site modelling, such as ASMC, for identifying isofunctional protein groups regardless of the taxonomic rank of the source organisms.

By grouping sequences according to the conservation of their genomic context, NetSyn appears to be a valuable tool for supporting the functional annotation of protein sequences with unknown function. Application of NetSyn to two protein datasets demonstrates its ability to gather enzymes acting in the same degrading system, even in the absence of sequence similarity. Their grouping is in fact due to the conservation of their genomic context which reflects underlying functional conservation. The integration of both homologous and non-homologous protein sequences as input is therefore essential when studying genomic structures such as operons, in which multiple genes involved in a specific cellular function (e.g., metabolic pathways, regulatory processes, etc.) are co-localized within the same locus. NetSyn is therefore an effective tool for highlighting groups of non-homologous proteins that participate in a common biological system, such as a metabolic pathway. Furthermore, we show that new putative enzymatic activities can be inferred from the annotation associated with genes in the genomic context of the target genes.

NetSyn is capable of constructing a network in which the relationship between two proteins depends only on the number of homologous genes within their respective genomic contexts. It can therefore be used either to explore the functional diversity of enzyme families or to identify conserved loci containing genes from different evolutionary origins involved in the same biological pathway (i.e., the same degradation/biosynthesis process). NetSyn thus addresses the challenging task of predicting novel enzymatic activities. NetSyn is freely available at https://github.com/labgem/netsyn.

In future developments, we aim to enable NetSyn to accept input files in GFF format, which would broaden its applicability and simplify the queries currently performed against UniProt and NCBI databases. Another improvement would be to incorporate additional sources of target proteins, such as NCBI GenPept, in order to mitigate the loss of diversity resulting from the removal of redundant proteomes in UniProt.

A further avenue for improvement would be to integrate gene transcription orientation into the computation of the synteny score. Currently, similar genomic contexts are identified solely based on the conservation of neighbouring genes, irrespective of their orientation. Conserved gene orientation could provide additional support for functional conservation when preserved. However, because supra-operonic structures (Pang and Lercher, 2017) are widespread in prokaryotic genomes and can maintain functional relationships despite variations in transcriptional orientation, gene orientation should be considered as complementary evidence rather than a strict criterion.

Finally, the node-merging functionality could be extended to automatically collapse redundant target proteins from closely related organisms sharing highly similar genomic contexts into a single representative prior to clustering. This preprocessing step would reduce the impact of taxonomic redundancy and promote functionally meaningful clustering

