## Supplemental data for "NetSyn: prokaryotic genomic context exploration of protein families"

### NetSyn supplementary data

##### S1 : Netsyn processing time

As NetSyn downloads the corresponding assembly files from the NCBI, the computation time for an analysis depends on server availability and load. Consequently, computation time may vary between analyses. Figure S1 illustrates the relationship between computation time and the number of input sequences from a dataset of 2,000 proteins of the xyloglucan degrading enzymes dataset. The computation was made on an intel Xeon E5-2697v4 cpu.

| Number of target protein sequences | Duration in minutes |
|---|---|
| 100 | 5 |
| 500 | 8 |
| 1000 | 11 |
| 2000 | 23 |

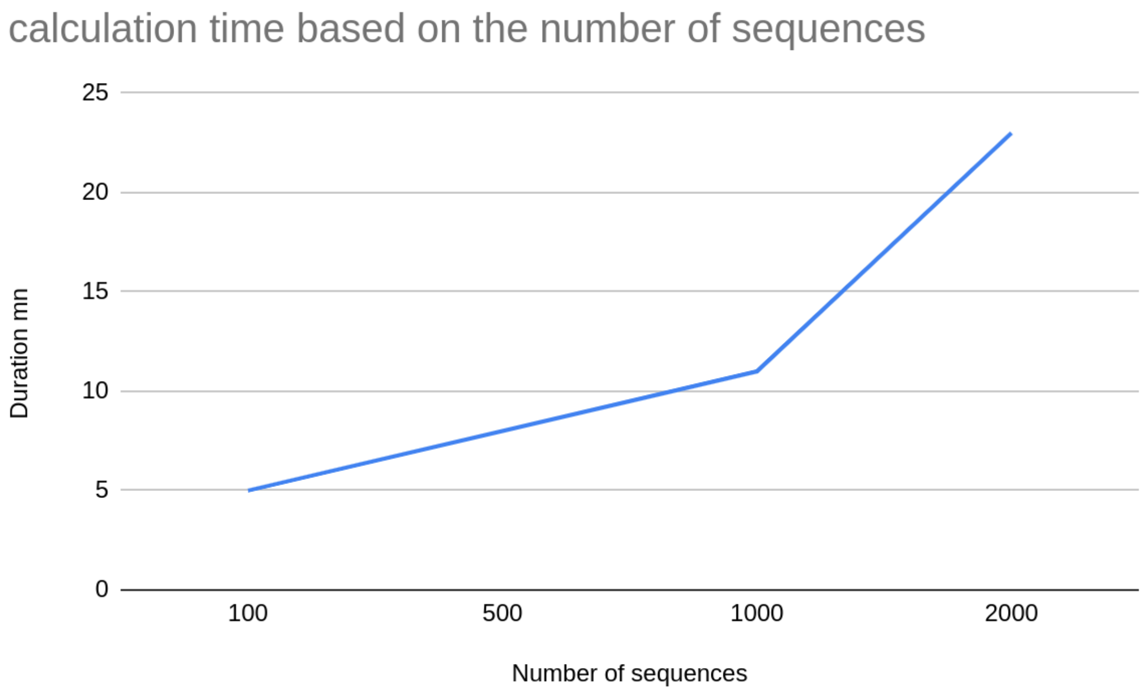

S2 : Example of a reducing network

Panel A shows a network with 438 nodes and 4334 edges.

Panle B shows the same network but nodes (i.e. proteins) belonging to a same netsyn cluster and coming from organisms belonging to the same order are merged into a same nodes. Therefore the reduced network contains only 117 nodes and 209 edges

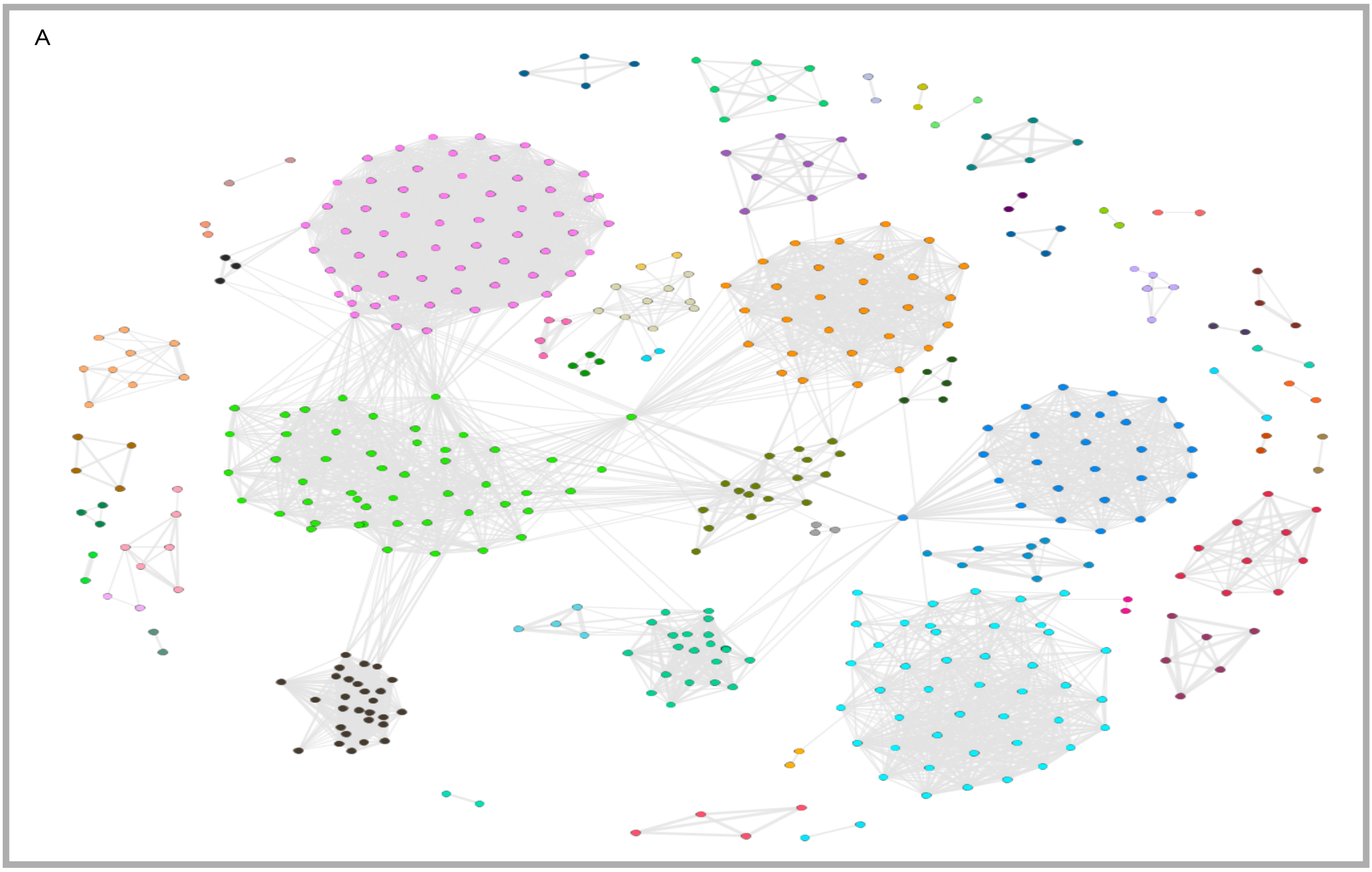


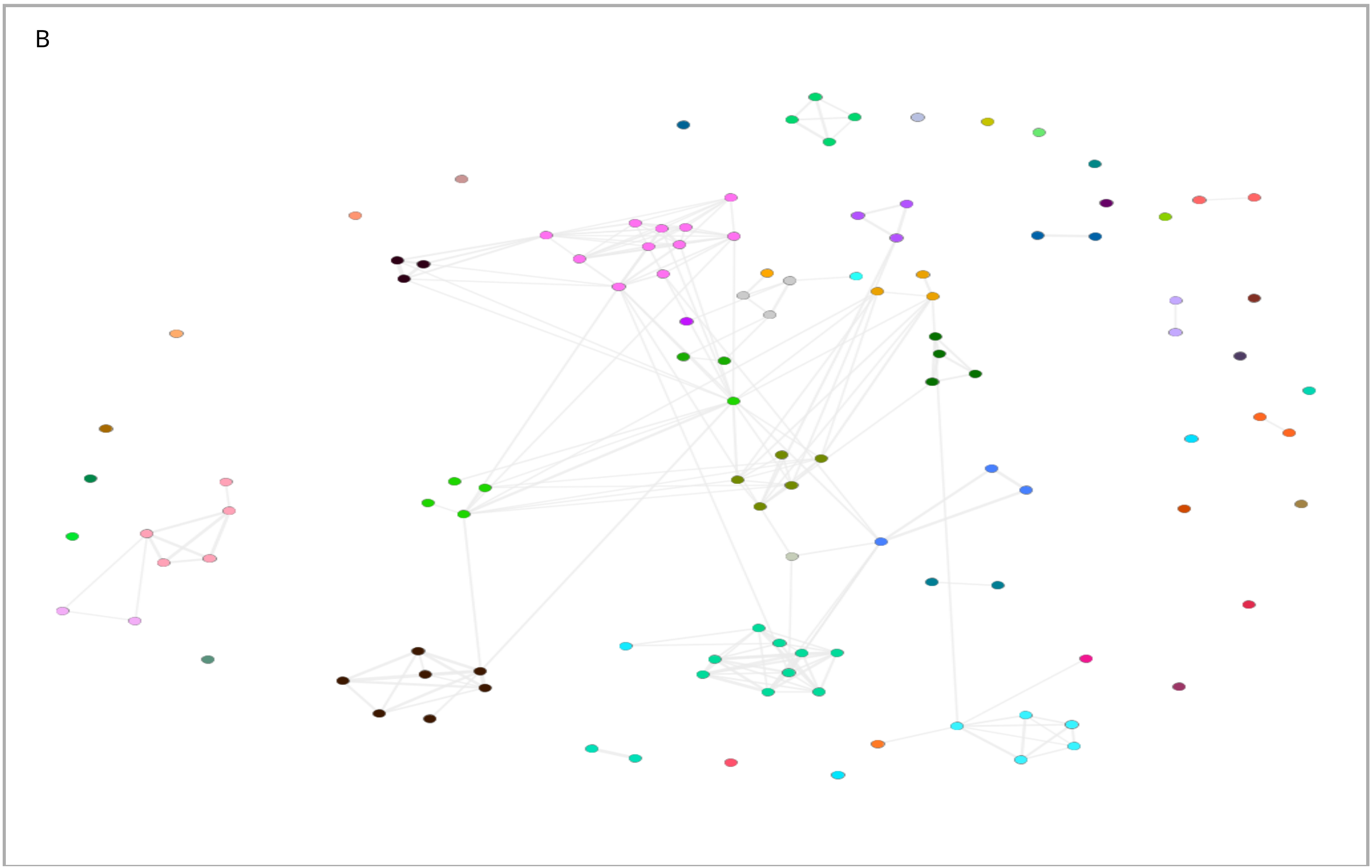

**S3: Genomic context of proteins belonging to NetSyn cluster of interest.** In bold are indicated enzymes which are specific to the pathway cited in the last column

| **ASMC Groups** | **NetSyn cluster (population)** | **Most popular genes in the neighborhood of BKACE genes as indicated in the NetSyn interface.** | **Prediction of pathway** |
|---|---|---|---|
| G1 | **Cluster 12**<br>(29 members from G1)<br>(2 members from G2)<br>(4 member from G3)<br>(4 members from G5)<br>(2 from G6)<br>(8 from G7)<br><br>Mostly alpha proteobacteria, beta-proteobacteria, small fraction of Chloroflexia and Chlorobia | amino acid transporter<br>oxidoreductase<br>3-hydroxybutyryl-CoA dehydrogenase<br>acetylpolyamine aminohydrolase<br>aminotransferase class III<br>acetyltransferase | **Conversion from valine to leucine: BKACE**<br>*[Monticello, D.J. & Costilow, R.N. Interconversion of valine and leucine by Clostridium sporogenes. J. Bacteriol. 152, 946–949 (1982).]* |
| G1 | **Cluster 22**<br>(32 members from G1)<br>(1 member from G4)<br>(1 member from G7)<br><br>(Mostly beta-proteobacteria and gamma-proteobacteria) | 3-ketoacyl-ACP oxidoreductase<br>GntR transcriptional regulator<br>MFS membrane protein transporter<br>cytosine deaminase<br>chloride channel<br>Di-heme cytochrome C peroxidase family<br>cytosine permease<br>short-chain dehydrogenase/reductase SDR | **Pathway to discover**<br><br>Could it be involved in the Catabolism of deoxyribose via glycerate and 2-phosophoglycerate ?<br><br>BKACE could cleave the beta keto acid from 2-deoxy-3-keto-ribonate (4,5-dihydroxy-β-ketopentanoate).<br><br>Protein C3K4S4 from Cluster 22 is homolog with 98% of sequence identity with A0A1N7U6D8, which correspond to sequence PS417_07250 described in [*Price M.N et al, Oxidative Pathways of Deoxyribose and Deoxyribonate Catabolism. mSystems. 4(1), 2019]* |
| G2 | **Cluster 19**<br>(27 members from G2)<br>(1 member from G1) | Zn-dependent alcohol dehydrogenase<br>**Lysine 2,3-aminomutase**<br>**L-beta-lysine 5,6 aminomutase**<br>3-oxoacid CoA transferase, alpha and beta subunit | **Lysine fermentation:**<br>BKACE catalyzes the cleavage of KAH in the context of lysine fermentation<br>*[Kreimeyer, A. et al. Identification of the last unknown genes in the fermentation pathway* |

| | | | |
|---|---|---|---|
| | (2 member from G3)<br>(1 member from G4)<br><br>Clostridia, Fusobacteriia, Bacteroidia, Beta-proteobacteria and Gamma-proteobacteria | B12 binding domain | *of lysine. J. Biol. Chem. 282, 7191–7197 (2007)]*<br>. |
| G2 | **Cluster 10**<br>(5 members from G2)<br>(2 membres from G7)<br><br>Only actinobacteria | aminomethyl transferase<br>Ferric uptake regulator<br>DUF1794<br>L-asparaginase<br>4-amino-4-deoxychorismate lyase | **Pathway to discover** |
| G3 | **Cluster 1**<br>(29 members from G3)<br>(3 members from G7)<br>(1 member from G1)<br><br>Alpha- and Beta-proteobacteria few Actinobacteria and 1 Thermoleophilia | Acyl-Coa dehydrogenase<br>Carboxyl transferase<br>Carbamoyl-phosphate synthase<br>Enoyl-CoA hydratase/isomerase<br>Long-chain-fatty-acid-CoA ligase<br>Short-chain dehydrogenase/reductase<br>glutathione-S-transferase<br>Fatty-acid desaturase<br>AraC transcriptional regulator | **Pathway to discover**<br>Probably Fatty acid biosynthesis |
| G4 | **Cluster 5**<br>(30 members from G5)<br>(1 member from G7)<br>Only alpha-proteobacteria) | **Protocatechuate 3,4-dioxygenase alpha and beta subunit**<br>**4-carboxymuconolactone decarboxylase/hydrolase**<br>4-hydroxybenzoate 3-monooxygenase<br>**Cis-muconate cycloisomerase**<br>transcriptional activator PcAQ<br>Glutathione-dependent formaldehyde dehydrogenase | **Catechol catabolism to beta-ketoadipate (pathway of aromatic compound degrada-<br>tion via protocatechuate to β-ketoadipate):**<br>BKACE metabolizes β-ketoadipate with the formation of acetoacetate<br>and succinyl-CoA<br><br>*[Collier, L.S., Gaines, G.L. III & Neidle, E.L. Regulation of benzoate<br>degradation in Acinetobacter sp. strain ADP1 by BenM, a LysR-type* |

| | | | |
|---|---|---|---|
| | | | *transcriptional activator. J. Bacteriol. 180, 2493–2501 (1998)]* |
| G4 | **Cluster 28**<br>(5 members) | membrane protein, ComEC/Rec2 family<br>alpha-ketoglutarate permease<br>chemotaxis sensory transducer<br>aldo/keto reductase | **Pathway to discover** |
| G4 | **Cluster 27**<br>(4 members) | ABC transporter<br>Cyclic nucleotide-binding, transcriptional regulator, Cpr/Fnr family<br>Cytochrome c class I<br>Uspa protein<br>Carbonate dehydratase<br>Dehydrogenase reductase | **Pathway to discover** |
| G5 | **Cluster 0**<br>(60 members)<br>(5 members from G1)<br>Mostly beta-proteobacteria<br>Actino-bacteria<br>Gamma-proteobacteria | **3-hydroxyacyl-CoA dehydrogenase**<br>glycine betain ABC transport<br>transcriptional regulator AraC<br>esterase<br>**Gamma-butyrobetaine dioxygenase** | **Carnitine degradation:**<br>BKACE catalyzes the condensation of dehydrocarnitine and acetyl-CoA into acetoacetate and betainyl-CoA [*Bastard and Smith, Nat. Chem. Biol. 2014*].<br><br>**Route 1:** conversion of γ-butyrobetaine into carnitine [*Uanschou, C., Frieht, R. & Pittner, F. What to learn from a comparative genomic sequence analysis of l-carnitine dehydrogenase. Monatsh. Chem. 136, 1365–1381 (2005).*] |
| G5 | **Cluster 7**<br>(47 members from G5)<br>(9 members from G1)<br>(4 members from G3)<br>(1 member from G4)<br><br>Mostly Bacilli And minority of gamma proteobacteria and delta proteobacteria | **carnitinyl-CoA dehydratase**<br>Transcription regulator AraC<br>**acyl-CoA dehydrogenase**<br>**acyl-CoA synthase**<br>3-ketoacyl-ACP reductase<br>aldo/keto reductase | **Carnitine degradation:**<br><br>BKACE catalyzes the condensation of dehydrocarnitine and acetyl-CoA into acetoacetate and betainyl-CoA [*Bastard and Smith, Nat. Chem. Biol. 2014*].<br><br>**Route 2: the first step uses three enzymes** (Acyl-CoA synthetase, Acyl-CoA dehydrogenase, carnitinyl dehydratase) instead of one (gamma-butyrobetaine dioxygenase) [*Kleber, H.P. Bacterial carnitine metabolism. FEMS Microbiol. Lett. 147, 1–9 (1997).*] |

| | | | |
|---|---|---|---|
| G6 | **Cluster 14**<br>(23 members from G6)<br><br>Alpha-proteobaceria<br>Gamma-proteobacteria<br>Beta-proteobacteria | **adenosylmethionine-8-amino-7-oxononanoate aminotransferase**<br>RpiR family,transcriptional regulator<br>TRAP transporter solute receptor<br>Universal stress protein UspA,<br>glutamate and aspartate transporter subunit | **Degradation of beta-aminotransferases**<br>BKACE decarboxylates Beta-ketoglutarate according to [ *Bastard and Smith, Nat. Chem. Biol. 2014*]. |
| G6 | **Cluster 8**<br>(7 members from G6) | **3-hydroxyacyl-CoA dehydrogenase**<br>LysR family regulatory protein<br>**Pyruvate carboxyltransferase**<br>**D-isomer specific 2-hydroxyacid dehydrogenase NAD-binding** | **Pathway to discover** |
| G7 | **Cluster 25**<br>(47 members from G7)<br>(7 members from G3)<br>(2 members from G2)<br>Mostly<br>Beta-proteobacteria<br>Actinobacteria<br>And a few gamma-proteobacteria | NADPH-containing alcohol dehydrogenase<br>**Alpha-ketoglutarate-dependent dioxygenase (TauD)**<br>MFS superfamily<br>DUF1234 (Serine hydrolase)<br>AraC transcriptional regulator<br>LuxR transcriptional regulator<br>acyl-CoA synthetase / fatty-acid ligase<br>**Thioredoxin / disulfide oxidoreductase** | **Pathway to discover**<br><br>Network described here:<br>https://string-db.org/cgi/network?taskId=bVB81MXU2ujy&sessionId=blujXK3InGmY |
| G7 | **Cluster 9**<br>(10 members from G7)<br>Only taxonomic order Burkholderiales | transcriptional regulator BetL<br>**betaine aldehyde dehydrogenase**<br>**choline dehydrogenase**<br>MFS sugar transport<br>rhamnosyltransferase | **Pathway to discover**<br><br>Could it be involved in the biosynthesis of the osmoprotectant glycine betaine ? [ Presence of a gene encoding choline sulfatase in *Sinorhizobium meliloti bet* operon: Choline-*O*-sulfate is metabolized into glycine betaine. Østerås, M *et al*, PNAS, 1998 95 (19) 11394-11399; |
| G7 | **Cluster 2**<br>(4 members from G7)<br>Mostly<br>Actinobacteria and<br>Deinococci | DUF985<br>Heat shock protein Hsp20<br>Glyoxalase resistance protein<br>Beta-ketoacyl synthase<br>DUF224 | **Pathway to discover** |

**S4:** Details of the active site logo of some enzymes that are gathered in the same NetSyn Cluster but belonging to different ASMC groups. Bold indicates the residues that are specific to the ASMC group. For G3, no specific residues exist but all catalytic residues are present.

| **Uniprot ID BKACE_id (from Bastard *et al.*, 2014)** | **ASMC group** | **NetSyn cluster** | **Annotation of common gene with G4 (ie NetSyn 19)** | **Quality of the model** | **Active site logo** |
|---|---|---|---|---|---|
| C6CDQ7 BKACE_759 | G3 | 19 | Acetoacetate:butyrate CoA-transferase (β subunit) Acetoacetate:butyrate CoA-transferase (α subunit) | 36% | AVHHTSGDSSDWFEFCV**I**RED |
| Q21BM8 BKACE_424 | G3 | 19 | Acetoacetate:butyrate CoA-transferase (β subunit) Acetoacetate:butyrate CoA-transferase (α subunit) | 34% | AMHHTSGDNSDWFEFCVVRED |
| A3XE58 BKACE_ 105 | G4 | 19 | Acetoacetate:butyrate CoA-transferase (β subunit) Acetoacetate:butyrate CoA-transferase (α subunit) | 79% | SLHHS-G-**R**SSFYEFVV**I**RED |
| Q216X8 BKACE_423 | G1 | 19 | Acetoacetate:butyrate CoA-transferase (β subunit) Acetoacetate:butyrate CoA-transferase (α subunit) | 54% | G**V**HHTGGLMSNFFEYV**I**ARED |

**S5 : presence or absence of co-localised key enzymes**

| organisms | GH31 | GH35 | GH95 | GH29 | GH74 |
|---|---|---|---|---|---|
| Xanthomonas_sp._WG16 | X | X | X | | X |
| Alteromonas_sp._RKMC-009 | X | X | X | | |
| Duganella_sp._leaf126 | X | X | X | | |
| Xanthomonas_arboricola_pv._pruni_MAFF_311562 | X | X | X | | X |
| Cellvibrio_sp._PSBB006 | X | X | X | | |
| Massilia_sp._CCM_9206 | X | X | X | | |
| Duganella_sp._Leaf61 | X | X | X | | |
| Xanthomonas_vasicola_CO-5 | X | X | X | | X |
| Pseudoxanthomonas_wuyuanensis_CGMCC_1.10978 | X | X | X | | X |
| Xanthomonas_pisi_LMG_847 | X | X | X | | X |
| Rhodanobacter_fulvus_Jip2 | X | X | X | | X |
| Xanthomonas_phaseoli_pv._syngonii_LMG_9055 | X | X | X | | X |
| Aestuariibacter_sp._GS-14 | X | X | X | | |
| Alteromonadaceae_bacterium | X | X | X | | |
| Pseudoduganella_albidiflava_KCTC_12343 | X | X | X | | X |
| Xanthomonas_sp._XNM01 | X | X | X | | X |
| Xanthomonas_fragariae_PD5205 | X | X | X | | |
| Massilia_timonae_NEU | X | X | X | | |
| Rhodanobacter_sp._Root627 | X | X | X | | X |
| Pseudoxanthomonas_suwonensis_11-1 | X | X | X | | X |
| Pseudoduganella_buxea_KCTC_52429 | X | X | X | | X |
| Xanthomonas_euroxanthea_CPBF_424 | X | X | X | | X |
| Duganella_sp._OV458 | X | X | X | | |
| Xanthomonas_hortorum_pv._carotae_CFBP_7900 | X | X | X | | X |
| Pseudoxanthomonas_sp._CF385 | X | X | X | | X |
| Xanthomonas_citri_pv._durantae_LMG696 | X | X | X | | X |
| Duganella_rivi_FT55W | X | X | X | | |
| Thermomonas_carbonis_KCTC_42013 | X | X | X | | X |
| Duganella_sp._CF402 | X | X | X | | |
| Duganella_vulcania_FT82W | X | X | X | | |
| Duganella | X | X | X | | |
| Xanthomonas_campestris_pv._campestris_B100 | X | X | X | | X |
| Xanthomonas_axonopodis_pv._melhusii_LMG9050 | X | X | X | | X |
| Massilia_aurea_CFS-1 | X | X | X | | |
| Duganella_margarita_FT134W | X | X | X | | |
| Dechloromonas_sp. | X | X | X | | X |
| Luteibacter_sp._OK325 | X | X | X | | X |
| Pseudoduganella_lutea_DSM_17473 | X | X | X | | X |
| Duganella_sp._SG902 | X | X | X | | |
| Xanthomonas_arboricola_pv._corylina | X | X | X | | |
| Xanthomonas_axonopodis_Xac29-1 | X | X | X | | |
| Cellvibrio_sp._KY-GH-1 | X | X | X | | |
| Xanthomonas_bromi | X | X | X | | |
| Pseudoduganella_namucuonensis_CGMCC_1.11014 | X | X | X | | |
| Xanthomonas_translucens_pv._poae_B99 | X | X | X | | |
| Marisediminitalea_aggregata_CGMCC_1.8995 | X | X | X | | |
| Xanthomonas_citri_pv._citri_JP541 | X | X | X | | X |
| Microbulbifer_rhizosphaerae_CECT_8799 | X | X | X | | X |
| Massilia_timonae_CCUG_45783 | X | X | X | | X |
| Pseudoxanthomonas_sp._3HH-4 | X | X | X | | |
| Stenotrophomonas_sp. | X | X | X | | |

| | | | | |
|---|---|---|---|---|
| Massilia_sp._Root335 | X | X | X | X |
| Microbulbifer_sp._SH-1 | X | X | X | |
| Pseudoxanthomonas_taiwanensis_J19 | X | X | X | |
| Xanthomonas_arboricola_Xanthomonas_arboricola_CFBP_8152 | X | X | X | X |
| Xanthomonas_campestris_pv._badrii_NEB122 | X | X | X | X |
| Xanthomonas_sp._Leaf148 | X | X | X | X |
| Xanthomonas_phaseoli_pv._phaseoli_CFBP6991 | X | X | X | X |
| Massilia_sp._YMA4 | X | X | X | |
| Massilia_sp._CCM_9029 | X | X | X | |
| Duganella_aquatilis_FT26W | X | X | X | |
| Massilia_oculi_CCUG_43427A | X | X | X | X |
| Xanthomonas_arboricola_pv._pruni_MAFF_301420 | X | X | X | X |
| Pseudoduganella_flava_CGMCC_1.10685 | X | X | X | X |
| Xanthomonas_arboricola_pv._guizotiae_CFBP_7409 | X | X | X | X |
| Xanthomonas_sp._3075 | X | X | X | X |
| Xanthomonas_campestris_pv._vesicatoria_str._85-10 | X | X | X | X |
| Teredinibacter_turnerae_T7902 | X | X | X | |
| Duganella_alba | X | X | X | |
| Massilia_cellulosiltytica_NEAU-DD11 | X | X | X | X |
| Saccharophagus_degradans_2-40 | X | X | X | |
| Duganella_sp._BJB489 | X | X | X | |
| Cellvibrio_japonicus_Ueda107 | X | X | X | |
| Luteimonas_salinisoli_SJ-92 | X | X | X | X |
| Pseudoxanthomonas_sp._Root630 | X | X | X | X |
| Luteimonas_wenzhouensis_YD-1 | X | X | X | |
| Xanthomonas_vesicatoria_ATCC_35937 | X | X | X | X |
| Xanthomonas_arboricola_pv._populi_CFBP3123 | X | X | X | X |
| Xanthomonas_sp._JAI131 | X | X | X | |
| Luteimonas_viscosa_XBU10 | X | X | X | X |
| Luteimonas_marina_FR1330 | X | X | X | X |
| Xanthomonas_campestris_pv._campestris_8004 | X | X | X | X |
| Xanthomonas_sp._ISO98C4 | X | X | X | X |
| Xanthomonas_axonopodis_DSM_3585 | X | X | X | |
| Pseudoduganella_umbonata_CECT_7753 | X | X | X | X |
| Xanthomonas_citri_pv._malvacearum_DSM_3849 | X | X | X | X |
| Duganella_flavida_FT135W | X | X | X | |
| _boreopolis_JCM_13306 | X | X | X | X |
| Duganella_sp._FT27W | X | X | X | |
| Massilia_aurea_AT3.2 | X | X | X | X |
| Duganella_violaceipulchra_HSC-15S17 | X | X | X | |
| Xanthomonas_vasicola_pv._vasculorum_SAM-118 | X | X | X | X |
| Xanthomonas_vesicatoria_53M | X | X | X | X |
| Pseudoxanthomonas_indica_P15 | X | X | X | |
| Xanthomonas_populi_LMG_5743 | X | X | X | |
| Xanthomonas_cucurbitae_CFBP2542 | X | X | X | X |
| Xanthomonas_citri_pv._eucalyptorum_LPF_602 | X | X | X | X |
| Salinimonas_sediminis_N102 | X | X | X | |
| Massilia_violaceinigra_B2 | X | X | X | |
| Xanthomonas_codiaei_CFBP4690 | X | X | X | X |
| Alteromonas_pelagimontana_5.12 | X | X | X | |
| Stenotrophomonas_sp._BIO128-Bstrain | X | X | X | X |
| Xanthomonas_arboricola_Xanthomonas_arboricola_CFBP_7645 | X | X | X | |
| Xanthomonas_citri_pv._vignicola_CFBP7111 | X | X | X | X |

| | | | | | |
|---|---|---|---|---|---|
| Pseudoxanthomonas_sp. | X | X | X | | X |
| Xanthomonas_hortorum_pv._gardneri_CFBP | X | X | X | | |
| Xanthomonas_sp._GW | X | X | X | | X |
| Duganella_radicis_KCTC_22382 | X | X | X | | |
| Duganella_vulcania_FT81W | X | X | X | | |
| Duganella_sp._BJB476 | X | X | X | | |
| Stenotrophomonas_panacihumi_JCM_16536 | X | X | X | | X |
| Xanthomonas_dyei_CFBP7245 | X | X | X | | X |
| Xanthomonas_prunicola_CIX249 | X | X | X | | X |
| Duganella_callida_DN04 | X | X | X | | |
| Massilia_sp._PDC64 | X | X | X | | X |
| Xanthomonas_campestris_pv._papavericola_NCPPB_2970 | X | X | X | | X |
| Xanthomonas_euroxanthea | X | X | X | | X |
| Xanthomonas_hortorum_pv._hederae_22-338 | X | X | X | | X |
| Duganella_guangzhouensis_FT80W | X | X | X | | |
| Xanthomonas_hortorum_Oregano_108 | X | X | X | | |
| Xanthomonas_phaseoli_pv._dieffenbachiae_LMG_25940 | X | X | X | | X |
| Xanthomonas_hortorum_pv._vitians_CFBP_498 | X | X | X | | |
| Massilia_sp._JS1662 | X | X | X | | X |
| Rheinheimera_sp._YQF-1 | X | X | X | | |
| Massilia_sp._CCM_9210 | X | X | X | | |
| Massilia_atriviolacea_SOD | X | X | X | | |
| Stenotrophomonas_terrae_DSM_18941 | X | X | X | | X |
| Xanthomonas_citri_pv._punicae_Xcp-4 | X | X | X | | X |
| Duganella_phyllosphaerae_T54 | X | X | X | | |
| Sphingobium | X | X | | X | |
| Duganella_vulcania_FT107W | X | X | X | | |
| Alteromonas_sp._38 | X | X | X | | |
| Xanthomonas_axonopodis_pv._vasculorum_NCPPB_900 | X | X | X | | |
| Cellvibrio_sp. | X | X | X | | |
| Xanthomonas_nasturtii_WHRI_8854 | X | X | X | | X |
| Pseudoduganella_plicata_KCTC_12344 | X | X | X | | X |
| Xanthomonas_axonopodis_pv._citri_306 | X | X | X | | X |
| Xanthomonas_hortorum_pv._pelargonii_CFBP_2533 | X | X | X | | |
| Cellvibrio_sp._79 | X | X | X | | |
| Pseudoxanthomonas_suwonensis_J1 | X | X | X | | X |
| Xanthomonas_hyacinthi_CFBP1156 | X | X | X | | |
| Parvularcula_dongshanensis_DSM_102850 | X | X | X | | X |
| Lysobacter_alkalisoli_SJ-36 | X | X | X | | |
| Pseudoxanthomonas_broegbernensis_NBRC_101013 | X | X | X | | X |
| Xanthomonas_floridensis_WHRI_8848 | X | X | X | | X |
| Massilia_arenae_GEM5 | X | X | X | | X |
| Xanthomonas_perforans_JK68-3 | X | X | X | | X |
| Massilia_forsythiae_GN2-R2 | X | X | X | | |
| Pseudoxanthomonas_sp._CF125 | X | X | X | | X |
| Microbulbifer_thermotolerans_DAU221 | X | X | X | | |
| Xanthomonas_sp._3793 | X | X | X | | |
| Lysobacter_penaei_SG-8 | X | X | X | | X |
| Xanthomonas_phaseoli_pv._phaseoli_CFBP6984 | X | X | X | | X |
| Pseudoduganella_dura_DSM_17513 | X | X | X | | X |
| Xanthomonas_prunicola_CFBP_8353 | X | X | X | | X |
| Xanthomonas_melonis_CFBP4644 | X | X | X | | X |
| Xanthomonas_arboricola_pv._populi_CFBP3123 | X | X | X | | X |

| | | | | | |
|---|---|---|---|---|---|
| Stenotrophomonas_rhizophila_BIGb0145 | X | X | X | | X |
| Xanthomonas_campestris_pv._campestris_ATCC_33913 | X | X | X | | X |
| Pseudoxanthomonas_sp._KAs_5_3 | X | X | X | | X |
| Xanthomonas_phaseoli_pv._manihotis_NBC1264 | X | X | X | | |
| Sphingopyxis_indica_DS15 | X | X | | X | |
